# Athletes exposed to uncommon vestibular stimulation strengthen their tactile-neural pathway

**DOI:** 10.64898/2026.05.05.722959

**Authors:** Rémi Demaria, Alix Moinon, Tom Negrel, Chloé Sutter, Jean Blouin, Martin Simoneau, Laurence Mouchnino

## Abstract

Strikingly, highly trained athletes engaged in vertiginous activities (e.g., *dance and slacklining*) and patients with bilateral vestibular loss show a similar pattern of neural plasticity, likely resulting from reduced vestibular sensory processes. However, unlike patients, these athletes show no balance impairments, quite the opposite. This suggests that the attenuation of vestibular processing represents an adaptive recalibration to excessive vestibular stimulation rather than a sign of dysfunction.

Concurrently, tactile processing increases as vestibular processing attenuates. Our findings indicate that effective adaptation extends beyond simple tactile compensation: it involves a strengthened tactile-brain pathway. Indeed, following unexpected base-of-support translations, the coupling between plantar shear forces (i.e., a proxy of plantar sole tactile afferents) and cortical responses over the somatosensory areas was markedly enhanced in Athletes. Cross-correlation analysis revealed stronger (r = 0.71) and faster (36 ms) tactile-brain coupling in Athletes (n = 25) compared with age- and gender-matched Controls (n = 18). This enhancement occurred within the first 180 ms following translation, that is, during the critical early phase of skin-surface interaction.

Notably, *artistic swimmers*, who undergo intense vestibular stimulation in a weightless underwater environment without balance equilibrium constraints, also exhibit enhanced tactile-brain coupling. This suggests that strengthening the tactile-brain coupling is not merely a byproduct of balance expertise, but rather a broader adaptive response to sustained vestibular stimulation. Multimodal neurons integrating vestibular and somatosensory inputs, such as those in the somatosensory cortex and thalamus, may increase their responsiveness to foot tactile afferents when vestibular inputs become excessive. In such contexts, the somatosensory system may assume a dominant role in providing gravity-related information for balance control.

## Introduction

Training involving extensive spinning and balancing modulates the sensory processing of vestibular afferents and promotes structural reorganization within the networks connected to the vestibular nuclei. This is demonstrated, for example, in highly trained dancers, ice dancing, and slackliners^1,2^. In these athletes trained in “vertiginous activities”, reduced volumes in the hippocampal formation, a key structure for spatial processing, have been observed^1^. Remarkably, a similar hippocampal volume reduction has been reported in patients with bilateral vestibular nerve section ^3^. Although comparable anatomical changes may arise from prolonged training or from vestibular loss, their behavioural consequences diverge substantially. On the one hand, in contrast to dancers, who demonstrate heightened awareness of their bodies and the surrounding space (e.g., 4, 5), patients with vestibular afferent deprivation show impairments in spatial tasks that require continuous updating of self-location and movement ^3^. On the other hand, vestibular patients typically exhibit compromised balance control ^6^, whereas dancers display precise dynamic equilibrium and postural control ^7^. The cerebellum, and specifically its regions encoding vestibular information, also undergoes transformation, resulting in reduced grey matter density, which is positively correlated with years of training in dancers ^2^. Overall, these cortical and cerebellar alterations suggest reduced vestibular afferent processing and improved filtering of irrelevant vestibular signals, which may help maintain sensorimotor performance during vertiginous activities.

Concomitantly, larger volumes in the posterior hippocampal formation and, of visual areas of the trained athletes were observed and interpreted as reflecting their increased reliance on visual cues to meet balance demands^1^. However, based on the seminal study of Fitzpatrick and McCloskey ^8^, the visual threshold for perceiving body movements was found to be higher than for somesthetic information. This means that, during standing, the receptors responsible for detecting early changes in body displacement relative to the supporting surface must be very rapidly adapting. This is because shear contact forces arising from skin-surface interaction have a dominant phasic component that requires rapid sensory encoding. Tactile receptors such as Pacinian corpuscles are consistent with this description ^9–11^. Even though athletes engaged in vertiginous activities are far from constituting a clinical population, they provide a valuable model for investigating downregulation of vestibular afferents, as their training exposes them to uncommon vestibular stimuli (i.e., large head acceleration).

Despite strong experimental evidence in patients with bilateral vestibular loss ^6,12^ and in monkeys ^13^ showing that somatosensory inputs are upregulated and can compensate for impaired vestibular-based graviception, it remains unclear whether these changes are restricted to higher-order cortical areas. A growing number of animal studies focused on somatosensory learning processes have revealed marked strengthening of information flow from tactile afferents to thalamo-cortical pathways ^14,15^. Therefore, it is reasonable to hypothesize that training-related changes are not confined to higher-order cortical areas but unfold dynamically across the entire pathway, from tactile afferent inputs to cortical processing.

This hypothesis is grounded in sensory weighting theory ^16,17^, which proposes that the central nervous system exerts dynamic control over afferent signal transmission and modulates cortical activity in a task-dependent fashion, beyond effects attributable to structural adaptations^18–21^. This task-dependent modulation is reflected in the inhibition of task-irrelevant cortical processes, as evidenced by alpha-band synchronization observed by Haufler et al.^22^ in the left central, temporal, and parietal regions of expert marksmen during the aiming phase of target shooting relative to novices (see 23, 24), for the gating role of alpha oscillatory activity).

Using early cortical responses to plantar tactile stimulation (SEPs), we should observe facilitation in Athletes engaged in vertiginous activities (dance and slacklining) compared with Control participants. This facilitation (e.g., greater amplitude and shorter latencies) is of particular interest because it reflects the earliest cortical processing of tactile afferents ^25–27^. Training-dependent strengthening in the tactile-brain coupling will be assessed by computing the cross-correlation between the lateral shear force (i.e., a proxy of plantar sole tactile afferents) and the evoked somatosensory signal recorded over the somatosensory areas. Finally, because tactile afferents can partially compensate for vestibular deficits^6^ by supporting the encoding of whole-body orientation relative to both the ground and the Earth’s vertical ^28,29^ behavioural responses to base-of-support translation are expected to be. shorter duration and greater magnitude (e.g., 30).

## Materials and methods

### EXPERIMENT 1

#### Participants and stimulating system

Forty-three participants performed the experiment. Among them, 25 participants were high-performing athletes (a mean of 5-9 hours of practice per week; an average of 12 years of practice) in vertiginous activities (12 dancers and 13 slackliners; hereafter referred to as the Athletes group), while the remaining 18 age- and gender-matched participants, who did not practice vertiginous activities, formed the Controls group. Participants reported being healthy (i.e., no neurological or musculoskeletal problems) and had comparable body mass index (BMI) values within the normal range (18.5 to 25 kg/m^2^). Participants read an information leaflet and signed an informed consent form before participating. This study followed the ethical standards of the latest version of the Declaration of Helsinki, except for database registration, and was approved by the research ethics committee (CERSTAPS: IRB00012476-2025-11-02-376).

Participants were asked to stand upright, barefoot, on a force platform, adopting a natural stance with the arms at the sides. Attention was paid to maintaining the self-selected foot position constant (i.e., feet shoulder-width apart before each trial) by marking foot placement on the force platform. A harness attached to a solid metal structure was secured to the participants’ waists as a safety measure to prevent falls. The force platform was mounted on rails (Bosh Rexroth) equipped with rolling to minimise friction. An electromagnet was installed on one side of the platform to keep it attached to the metal structure. Deactivating the electromagnet enabled platform acceleration (Fig. 1A). The load (∼2% of body weight) was adjusted to each participant’s weight, to ensure that switching off the electromagnet generated a standardized rightward platform acceleration of 33 ± 4 cm/s² (Fig. 1B).

**Figure 1.**
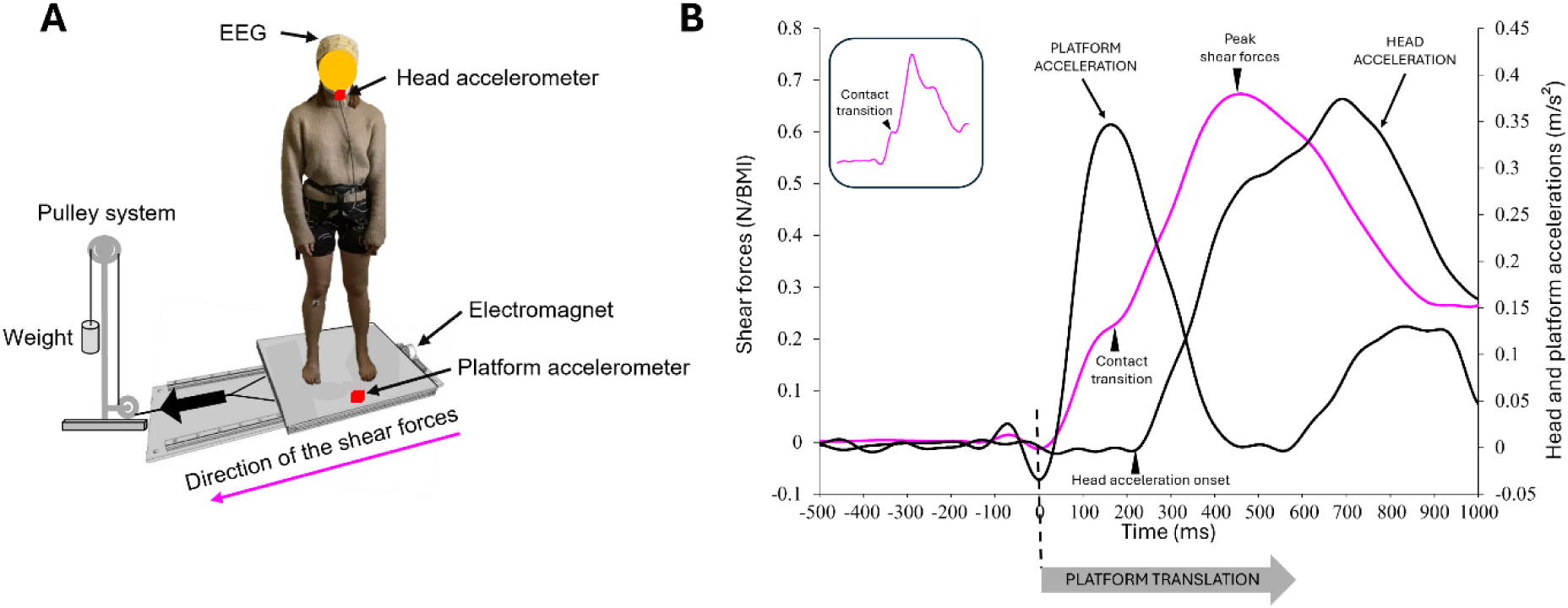
Experimental setup. **A.** Participant stood barefoot on a force platform, and electromagnet deactivation caused platform translation. **B.** One participant’s trial-average (n = 37) of shear force (N/BMI), head and platform accelerations (m/s²) as a function of time. The signals are plotted together to illustrate their temporal relationships. Time 0 (black broken line) indicates platform translation onset, plantar skin and platform surface interaction. The insert represents a single trial for the same participant.

Participants were asked to close their eyes upon receiving the verbal warning indicating that the trial was about to begin. In all trials, participants had to maintain an upright, steady posture for 7 s (i.e., the duration of the trial recording). To prevent participants from anticipating the platform acceleration onset, the platform translation occurred at any time between 2 and 4 s after the verbal signal. In addition, we included eight trials without any platform acceleration. These trials were randomly distributed among the 37 trials with platform acceleration, with the same trial order presented to all participants. Short breaks were frequently offered to participants during the experiment to reduce fatigue.

Before the experimental session, the Semmes-Weinstein monofilament test ^31^ was used to assess plantar sole sensitivity to static mechanical stimuli. Each monofilament had a nylon thread calibrated to a specific weight (in grams) at its tip. The test was carried out on each plantar sole on two areas, with a relatively high density of mechanoreceptors ^32^. Pressure was applied perpendicular to the skin surface, and participants, with their eyes closed, were asked to report whether they perceived it.

#### Behavioural recordings and analyses

The ground reaction forces and moments were recorded using an AMTI force platform (60 × 120 cm, Advanced Mechanical Technology Inc., United States) at a sampling frequency of 1000 Hz and filtered offline using a dual-pass 4th-order Butterworth low-pass filter with 10 Hz cut-off frequency. The signals were then rebased using a one-second recording period preceding the onset of the platform acceleration. Shear forces were analysed only along the mediolateral (ML) axis, as force changes in this direction represent the earliest signature of cutaneous stimulation during platform acceleration. The shear forces were normalised relative to the participant’s BMI. We defined the shear force onset when the force started to increase consistently (Fig. 1B). The shear forces increased until a transition phase, where the forces briefly decreased. This sharp force change, previously reported in studies using similar platform translations (e.g., 33, 34), was evident in individual trials (see inset in Figure 1B), but was smoothed out by temporal variability when averaging all trials. We defined the period between the onset of shear force and its transient decrease as the “*early phase*”. The force amplitude at the end of that phase was quantified relative to a 1-s baseline period computed prior to the translation. Due to the body’s inertia, the platform displacement is accommodated primarily by skin deformation, leading to relative sliding or friction between contacting surfaces (i.e., foot skin-surface interaction). Afterwards, the shear forces continued to increase until a peak was reached before reversing their direction. The period between the end of the early phase and this peak force is defined as the “*late phase*” and is considered a postural response aimed at correcting body sway.

We measured head acceleration (Fig. 1B) to estimate the timing and magnitude of vestibular stimulation induced by the platform translation. Head acceleration was measured with a triaxial accelerometer (model 4630, Measurement Specialties, USA) mounted on the participants’ chins. The acceleration signals were recorded at 1000 Hz and then processed offline with a dual-pass 4th-order Butterworth low-pass filter with a cutoff frequency of 10 Hz. Based on Gianna et al.^35^, the vestibular stimulation was assumed to occur when head lateral acceleration exceeded a threshold of 0.048 m/s². Determining the onset of vestibular stimulation was crucial for establishing when head motion occurred relative to tactile stimulation, allowing us to distinguish cortical responses driven by plantar sole afferents from those driven by vestibular afferents.

##### Electroencephalography recordings and analyses

Electroencephalographic (EEG) activity was recorded continuously from a 64 Ag/AgCl surface-electrode elastic cap, sized to the participants’ heads (ActiveTwo system, BioSemi, Amsterdam, The Netherlands). The system included two specific electrodes: an active CMS (Common Mode Sense) electrode and a passive DRL (Driven Right Leg) electrode, replacing the conventional reference electrode. In this configuration, the CMS electrode detected the signal common to the electrodes, while the DRL electrode reduced noise. This CMS/DRL loop therefore stabilised the EEG signal. The signals were pre-amplified at the electrode sites, post-amplified with DC amplifiers, and digitized at a sampling rate of 1024 Hz. The signals were further resampled at 1000 Hz and, if necessary, filtered off-line with a 50 Hz digital notch filter (24 dB/octave), implemented in BrainVision Analyser 2 software (Brain Products, Germany). After artefact rejections based on visual inspection, 98% of trials were included in the analyses.

Somatosensory evoked potentials (SEP) were obtained by averaging all epochs time-locked to the shear force onset for each participant. The 50 ms interval preceding support translation served as the baseline for SEP averaging. Consistent with prior studies using lower-limb stimulation (e.g., 33, 36, 37), SEP amplitudes were maximal at the Cz electrode. This electrode overlies the sensorimotor cortices, where the foot representation is located along the medial wall within the longitudinal fissure, as depicted in the cortical homunculus. Analyses focused primarily on the P1N1 wave following sensory stimulation and, secondarily, on the subsequent P2N2 component. For each component, peak-to-peak amplitude and component latencies were analysed.

##### Cross-correlation analysis

To assess the strength of coupling between sensory afferents and cortical responses, we computed the cross-correlation between the shear forces (i.e., a proxy of plantar sole afferents) and the evoked cortical responses over the somatosensory areas on individual trials (n = 37). Cross-correlation assumes a linear relationship between the variables being cross-correlated. The cortical signals were shifted in time relative to the shear force signal over a 300-ms time window during the early and the late phases. The correlation coefficient was computed between the two time-series using a 1-ms step. The time lag associated with the maximum correlation coefficient was used to determine whether, and by how much, the cortical signal led (negative value) or lagged (positive value) the force signal. The cross-correlation analysis was computed using the MATLAB cross-correlation (*xcorr*) function (Version R2024b, The MathWorks Inc., Natick, United States).

### EXPERIMENT 2

To further determine whether changes in plantar sole tactile-cortical coupling or SEPs are driven by balance-control expertise in dancers and slackliners rather than by extensive vestibular stimulation, we tested a group of artistic swimmers. These Athletes are exposed to substantial vestibular stimulation, often involving rapid head rotations in inverted positions, but without comparable balance-equilibrium constraints. Indeed, their training is performed predominantly in non-weight-bearing conditions. Comparing this group with dancers and slackliners, therefore, allows us to dissociate the effects of balance-control expertise from those of vestibular stimulation per se. Twelve participants (12 females; age: 21 ± 1 years; height: 164 ± 7 cm; weight: 59 ± 5 kg; BMI: 22 ± 2 kg/m²), who practised artistic swimming (a mean of five training sessions per week) followed the same protocol as in Experiment 1. The procedure and analyses were identical to those of Experiment 1.

### EXPERIMENT 3

To identify the origin of the cortical SEP changes, direct recordings were made before spinal cord relay. In a third experiment, the transmission and processing of tactile receptors was assessed by measuring the response evoked by the electrical stimulation of the participants’ right mixed-tibial nerve (i.e., somatosensory and motor). We recorded peripheral nerve action potentials at the popliteal fossa just above the fossa crease (see 38, for review) while participants maintained upright and stable position. Ten participants (women) different from experiments 1 and 2, took part in this experiment: 5 Artistic swimmers (mean age 22 ± 2 years; mean weight 61 ± 5 kg; mean height 168 ± 7 cm) and 5 Controls (mean age 23 ± 2 years; mean weight 59 ± 5 kg; mean height 167 ± 6 cm). Fifteen stimulations were delivered via two electrodes (Fyzea Optimum Electrodes) by an isolated bipolar constant current stimulator (DS5 Digitimer, Welwyn Garden City, UK) over the 10 recordings of 15 seconds duration, giving a total of 150 stimulations. The electrodes were located on the medial malleolus ^39^ (Fig. 2A). The inter-stimulus interval was randomly chosen between 800 and 1200 ms to limit participant’s anticipation of the next stimulation. Because the inter-stimulus intervals were > 800 ms, the evoked response was unlikely influenced by the previous stimulation. When the inter-stimulus intervals were < 500 ms, a decrease in the evoked response was commonly observed due to the previous stimulation ^40^. For each participant, the current used to stimulate the tibial nerve (mean: 2 ± 0.4 mA) was set 25% above the perceptual threshold but remained below the threshold for evoking motor movements. This threshold was estimated before the experimental session while the participants were standing upright and did not differ between the two groups (F_1,8_ = 1.56; *p* = 2.46).

**Figure 2.**
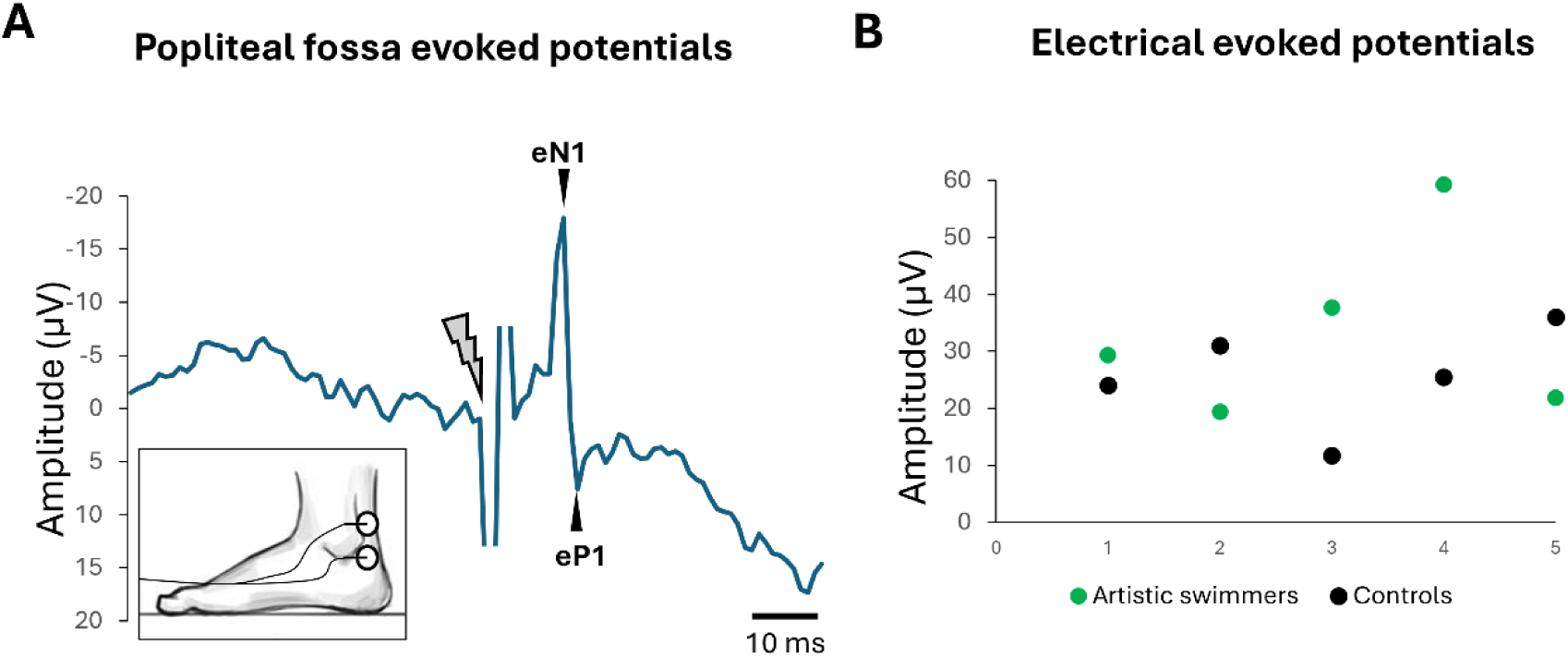
**A.** Averaged (n = 150) tibial nerve evoked potential recorded at the popliteal fossa for one participant. The broken lines correspond to the stimulation artifact. The insert represents the electrodes placement. **B**. Mean amplitude of the evoked potentials for each participant.

#### Statistics

For each dependent variable, we assessed homogeneity of variances using Levene’s test and normality using the Kolmogorov-Smirnov test. The significance threshold was set at p ≤ 0.05. To assess the strength of the coupling between shear force and somatosensory evoked potentials, we compared the mean values for Athletes, Controls (Experiment 1), and Artistic swimmers (Experiment 2 and 3) using separate repeated-measures one-way ANOVAs. Furthermore, we used paired t-tests to compare P1 and N1 latencies relative to the time point at which the interaural head acceleration exceeded the vestibular threshold. Effects that showed statistical significance were subjected to a Newman-Keuls post hoc test for further comparison. As the results of the Semmes-Weinstein monofilament test were not normally distributed, we used non-parametric Chi-square (χ²) and Kruskal-Wallis *H* tests.

## Results of Experiment 1

### Plantar Tactile Sensitivity

We first examined whether plantar sole tactile sensitivity thresholds varied across groups. Since no differences were observed across the 4 tested sites (first toe and medial metatarsals of both feet) (χ² = 3.15; p = 0.36 and χ² = 1.13; p = 0.76, for Controls and Athletes respectively), we averaged the 4 measurements for each participant. The results did not show a significant difference in the overall tactile sensitivity threshold between the two groups (KW-*H_1_*_,41_ = 0.53; *p* = 0.47; Fig. 3A). The mean plantar sole sensitivity threshold was 0.44 grams, which is within the range of healthy adults ^41^. Thus, any differences observed between Athletes and Controls in behavioural or cortical responses cannot be attributed to differences in plantar sole sensitivity, at least for the indentation levels tested with monofilaments.

**Figure 3.**
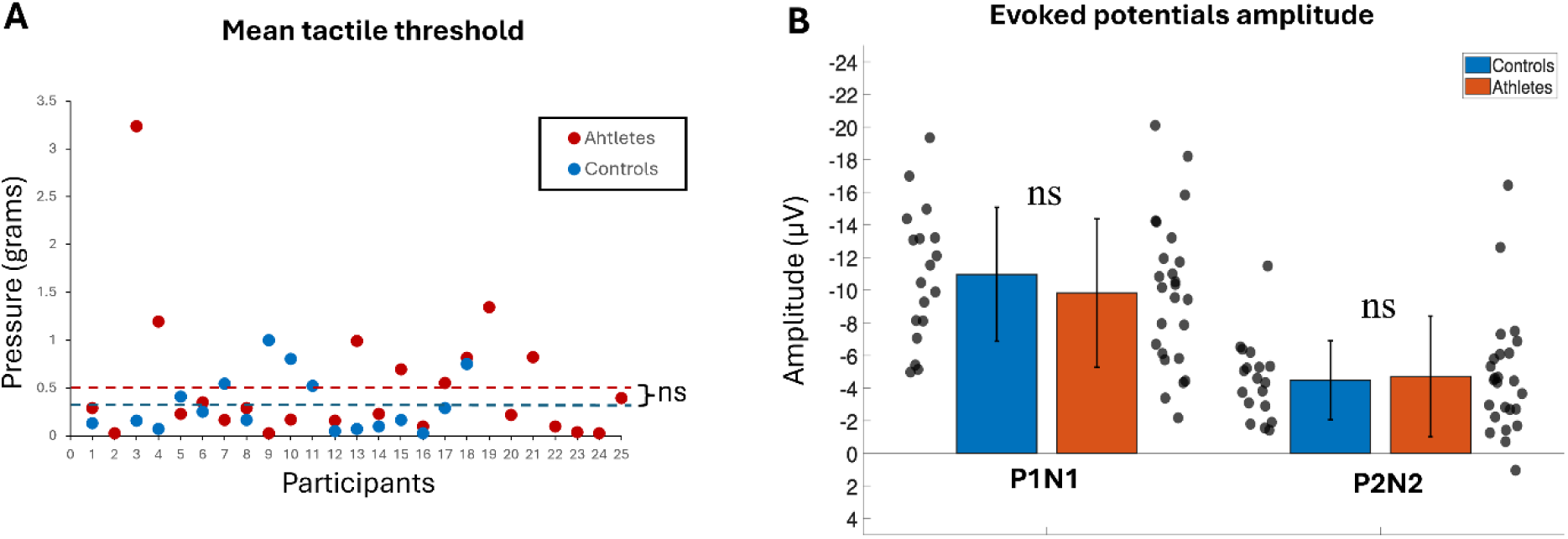
**A.** Mean pressure of the plantar tactile threshold for the Control and Athlete groups. Each dot represents a participant’s mean value. The dashed horizontal lines indicate the mean value for each group (ns, not significant). **B**. Mean amplitude of P1N1 and P2N2 SEPs. Error bars represent the group standard deviation.

### Cortical responses to sensory stimulation

In our previous work, participants with deprived vestibular afferents showed a larger P1N1 SEP in response to plantar tactile stimulation while standing ^12^. Here, we tested whether this somatosensory facilitation also occurs in Athletes performing vertiginous activities. Contrary to our hypothesis, P1N1 amplitude did not differ significantly between groups (*F*_1,41_ = 0.70; *p* = 0.41, overall mean of 10.30 ± 4.35 µV; Fig. 3B). Following the P1N1 complex, a distinct second component (P2N2) emerged over the somatosensory cortices at approximately half the amplitude of P1N1. Like P1N1, P2N2 amplitude did not differ significantly between Athletes and Controls (*F*_1,41_ = 0.05; *p* = 0.82; overall mean of 4.61 ± 3.19 µV; Fig. 3B).

In addition, no significant group effect was observed on P1 latencies (*F*_1,41_ = 1.97; *p* = 0.17; overall mean latency of 158 ± 25 ms). Interestingly, N1 latency occurred earlier in Athletes (225 ± 26 ms) than in Controls (243 ± 31 ms) groups (*F*_1,41_ = 4.08; *p* = 0.05; Fig. 4B). Consistent with this earlier N1, P2 and N2 latencies were also shorter in Athletes compared to Controls (*F*_1,41_ = 9.10; *p* = 0.004 and *F*_1,41_ = 7.26; *p* = 0.01 for P2 and N2, respectively; see Table 1). This P2N2 complex likely corresponds to a multisensory integration process combining at least tactile and vestibular afferents as the head is starting to move (see Table 1; 33).

**Figure 4.**
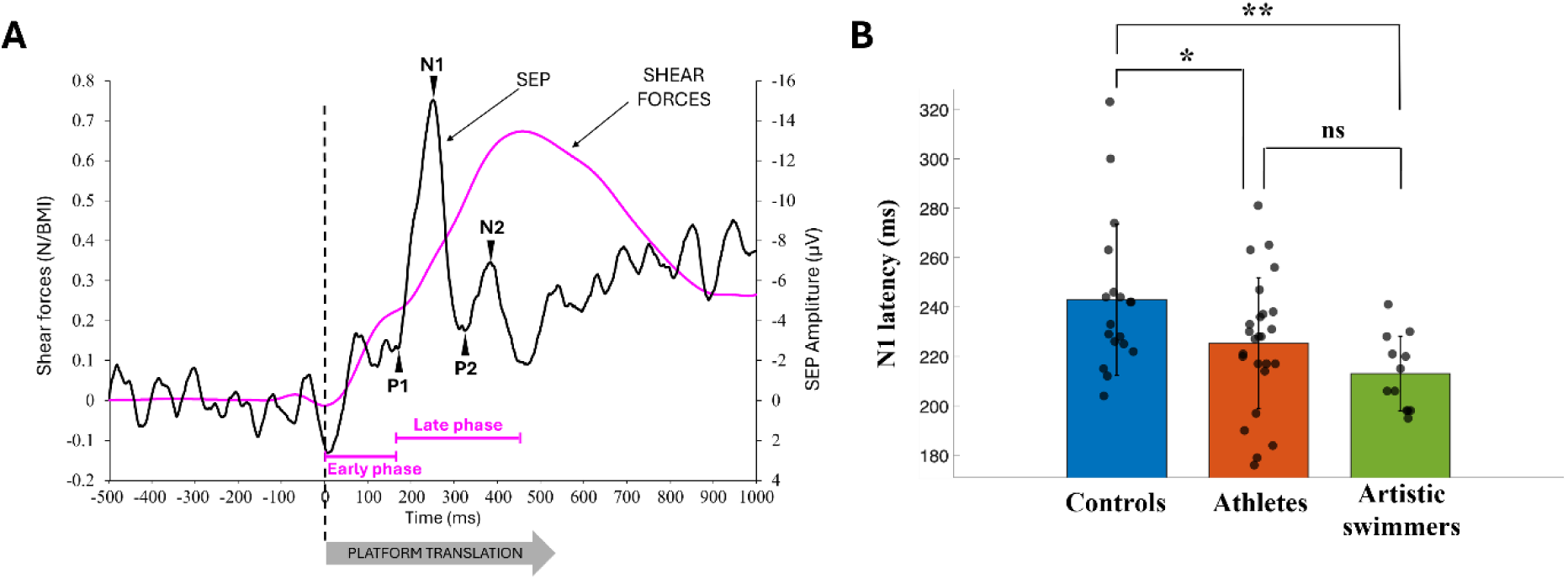
The black curve represents the SEP (Cz electrode) trial-average of one participant. The pink curve shows the average shear force for this participant’s trial. The dashed vertical line indicates the onset of platform translation. **B.** Group means N1 latency. Each dot represents a participant’s mean value. Error bars represent the group standard deviation (ns, not significant; *, p < 0.05; **, p < 0.01).

**Table 1.** The paired t-test compares P1, N1, and the vestibular threshold for Athletes and Controls (± SD).

| Controls |  |  |  | Athletes |  |  |  |
| --- | --- | --- | --- | --- | --- | --- | --- |
| Vestibular threshold |  |  |  |  |  |  |  |
| 279 ± 53 ms |  |  |  | 256 ± 37 ms |  |  |  |
| P1 | N1 | P2 | N2 | P1 | N1 | P2 | N2 |
| 164 | 243 | 321 | 375 | 153 | 225 | 288 | 343 |
| ± 19 ms | ± 31 ms | ± 38 ms | ± 38 ms | ± 27 ms | ± 26 ms | ± 33 ms | ± 38 ms |
| t-test |  |  |  |  |  |  |  |
| $t = 10.97$<br>$p \leq 0.001$ | $t = 4.16$<br>$p \leq 0.001$ | | | $t = 11.94$<br>$p \leq 0.001$ | $t = 3.54$<br>$p = 0.002$ | | |

On average, head acceleration exceeded the vestibular threshold 279 ± 53 ms and 256 ± 37 ms after platform translation onset in the Control and Athlete groups, respectively (no group difference: *F*_1,40_ = 2.80; *p* = 0.10). This long delay between platform translation onset and vestibular stimulation (i.e., head motion) onset can be attributed to whole-body inertia, which resists motion and thereby delays head and whole-body movements. Paired t-tests (see Table 1) showed that both P1 and N1 latencies significantly preceded vestibular stimulation, with mean lead times of 108 ± 43 ms for P1 and 33 ± 40 ms for N1. No group differences were observed in the lags between vestibular stimulation and either P1 or N1 (*F*_1,40_ = 0.94; *p* = 0.34 and *F*_1,40_ = 0.21; *p* = 0.65 for P1 and N1 components, respectively). These results suggest that the P1N1 cortical complex was unlikely evoked by head motion and was instead driven by plantar sole tactile afferents arising from the early foot skin stretch at platform acceleration onset.

### Behavioural responses to platform translation

The *early phase* of shear forces did not show differences in duration or amplitude between the Athlete and Control groups (F_1,41_ = 3.04; p = 0.09; and F_1,41_ = 1.92; p = 0.17 for duration and amplitude, respectively). The *early phase* lasted, on average, 179 ± 22 ms and reached a mean force amplitude of 5.6 ± 0.8 N (0.26 ± 0.04 when normalized to body mass index, N/BMI). This phase is thought to arise passively from friction between the stretched skin of the foot’s sole and the platform, and previous studies using the same platform acceleration have shown that it is not associated with changes in muscular activity^34^.

The *late phase* did not differ between the Athlete and Control groups in either duration (271 ± 55 ms) or amplitude (9.5 ± 1.8 N; 0.45 ± 0.09 N/BMI) (*F*_1,41_ = 0.17; *p* = 0.68 and *F*_1,41_ = 0.13; *p* = 0.72 for duration and amplitude, respectively). In contrast to the early phase, Sutter et al.^34^ demonstrated that the shear forces observed during the *late phase* were actively generated by participants and correspond to a balance-correcting response.

### Coupling between sensory inputs and cortical response

Next, we assessed the relationship between plantar sole tactile afferents and cortical response by computing the cross-correlation between the shear forces and the EEG signal recorded at the Cz electrode for individual trials. We observed a strong correlation (i.e., over 0.65) in both groups during the *early phase* and *late phase* (Fig. 5A). During the *early phase*, a main effect of group (*F*_1,41_ = 5.39; *p* = 0.03) confirmed stronger coupling in Athletes (0.71 ± 0.08) than in Controls (0.65 ± 0.07). The analysis revealed no group difference for the *late phase* (*F*_1,41_ = 0.94; *p* = 0.34; overall mean: 0.74 ± 0.08).

**Figure 5.**
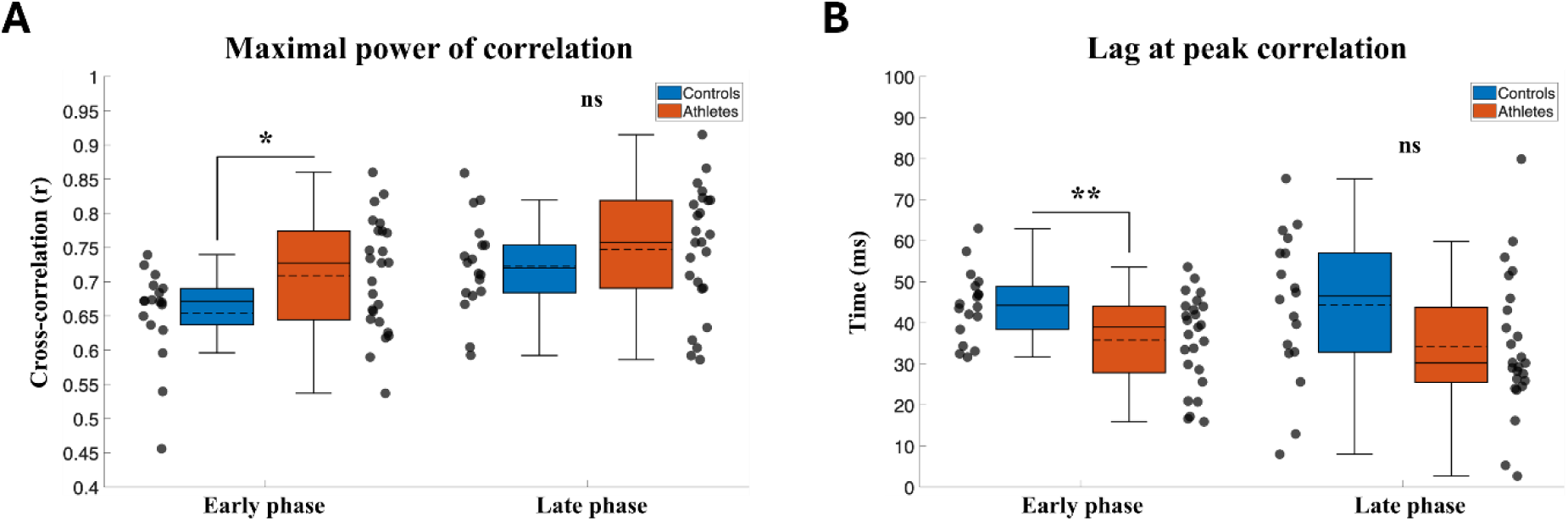
**A.** Group means for the coefficient of correlation. Each dot represents a participant’s mean value. **B.** Group means for the time lag at peak correlation. Error bars depict group standard deviation, and ns stands for not significant difference, while one (*) and two (**) asterisks indicate a significant difference with p < 0.05 and p < 0.01, respectively.

As expected, the cortical signals lagged the force signals (Fig. 5B). During the *early phase,* the correlation peak occurred earlier in Athletes (36 ± 11 ms) than in Controls (44 ± 8 ms) (group main effect: *F*_1,41_ = 7.34; *p* = 0.01), while no main effect of group was observed during the *late phase* (*F*_1,41_ = 3.61; *p* = 0.06; overall mean: 38 ± 18 ms).

The higher correlation peak and shorter delay observed in Athletes compared to Controls during the early phase suggest greater neural efficiency in processing tactile plantar sole afferents. During this early phase, the absence of head motion acceleration suggests that vestibular afferents did not contribute to strengthening the pathway between sensory afferents and cortical activity.

## Results of Experiment 2

Including a group of artistic swimmers who, unlike dancers and slackliners, train exclusively in water without foot contact with a solid surface and are routinely exposed to inverted positions and body-spinning manoeuvres, allows us to disentangle adaptations arising from foot-based balance training from those driven by excessive vestibular stimulation (i.e., uncommon head acceleration).

We compared the maximal coupling between shear force and evoked cortical responses across 3 groups: Artistic swimmers, Athletes, and Controls (the latter two from Experiment 1). The ANOVA revealed a main effect of Group, with stronger coupling (*F*_2,52_ = 3.89; *p* = 0.026) in artistic swimmers (0.72 ± 0.05) than in Controls (0.65 ± 0.07). No difference was observed between the Artistic swimmer and the Athlete groups (*p* = 0.77; overall mean: 0.71 ± 0.07; Fig. 6A). During the *early phase*, the analysis revealed group differences for the lag of the maximal coupling between the cortical and force signals (*F*_2,52_ = 12.56; *p* < 0.001). The lag occurred earlier in the Artistic swimmers (27 ± 5 ms) than in both Controls (44 ± 8 ms; p < 0.001) and Athletes (36 ± 11 ms; *p* = 0.008, Fig. 6B).

**Figure 6.**
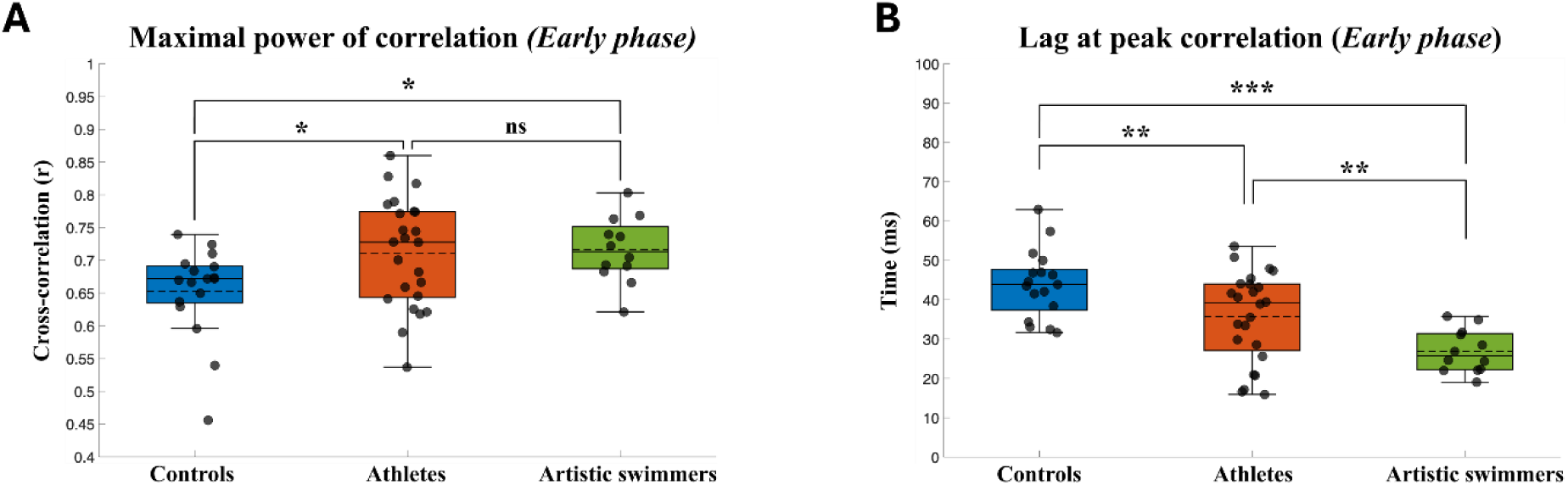
**A.** Group means for peak correlation (i.e., maximal power of correlation). Each dot represents a participant’s mean value. **B.** Group means for the lag at peak correlation. Error bars depict the group standard deviation, ns stands for not significant, while asterisks indicate group differences (*, *p* < 0.05, ** p < 0.01, and ***, p < 0.001).

The P1N1 SEPs amplitude in Artistic swimmers did not differ from that of Controls or Athletes (*F*_2,52_ N= 0.37, *p* = 0.70; overall mean: 10.25 ± 4.32 µV), and no differences were observed for P1 latency (*F*_2,52_ = 1.29; *p* = 0.28; overall mean:157 ± 22 ms). In contrast, N1 latency was shorter in Artistic swimmers than in Controls (*F*_2,52_ = 5.08; *p* = 0.005), while it remained similar between Artistic swimmers and Athletes (*p* = 0.18; overall mean of 221 ± 24 ms) (Fig. 4B).

## Results of Experiment 3

During quiet standing without any expected platform translation, the foot electrical stimulation evoked consistent responses for all participants (Fig. 2A). To assess whether training in vertiginous activity altered the transmission of tactile afferents at the very early stage of transmission, we compared the amplitude of the electrically evoked potentials between Artistic swimmers and Controls. Figure 2B depicts, for each participant, the differences in the amplitudes of the eN1-eP1. We observed no significant difference between groups in the evoked eN1eP1 amplitude (F_1,8_ = 0.89; *p* = 0.37), suggesting that distal nerve conduction was not affected by the specific training. This finding further indicates that the tactile P1N1 facilitation observed at the cortical level in Experiment 2 likely did not originate from peripheral mechanisms or modulation occurring prior travelling up to the spinal cord.

## Discussion

Considering the vestibular downregulation of Athletes involved in vertiginous activities (dancers and slackliners) and the significant sensorimotor balance requirements associated with these activities, we hypothesized that tactile afferent processing would be facilitated and tactile-brain coupling strengthened to improve motor responses during balance recovery. Closely matching our predictions, the results showed that, compared to controls, athletes showed: (1) shorter latencies of the somatosensory cortical potentials evoked by support platform translations, (2) greater tactile-brain coupling, and (3) shorter lag of the tactile-brain coupling, without changes in balance-correcting response. These findings held even when considering Artistic swimmers who experience prolonged training in the absence of equilibrium constraints while being exposed to extensive vestibular stimulation (i.e., head spinning).

The strengthening of the tactile-brain coupling occurred during the *early phase* of shear forces induced by platform translation. This phase primarily reflects the mechanical interaction between the plantar skin and the support surface, which generates tactile cues. Specifically, the Pacini corpuscles are particularly well suited to detect the friction induced vibrations generated by the relative motion between the foot’s skin and the contacted surface (e.g., 42). It is noteworthy that SEP components scaled systematically with vibration amplitude^43^. These mechanoreceptors embedded in the foot sole form a complex network, with particularly high density in the metatarsophalangeal region^44^. This density is especially important for balance control, as the forefoot bears more than 80% of the body’s weight ^45^.

The stronger coupling observed in Athletes, together with its shorter lag (from 27 to 36 ms), between shear forces (i.e., tactile inputs) and somatosensory activity may also be expressed at earlier processing stages of the ascending tactile sensory pathway. Because no change was observed (Experiment 3) at the level of peripheral nerve it seems unlikely that the facilitation takes place at this early stage. Travelling up to the cuneate nucleus, a lower brainstem structure that receives monosynaptic inputs from the primary afferents innerving the skin mechanoreceptors ^46^ with short latencies (12.5 ± 1 ms in the cat ^47^), this pathway could be part of this facilitated transmission.

This facilitation could also involve thalamocortical circuits, including the ventral posterolateral (VPL) nucleus, which exhibit increased excitatory synaptic strength following somatosensory learning ^14^ and respond to vibrotactile stimulation within tens of milliseconds ^48^. Moreover, the observed coupling strengthening could be further shaped by descending cortical projections: sensorimotor cortex sends cortico-cuneate projections that modulate tactile transmission in the cuneate nucleus ^49,50^, and corticothalamic feedback from S1 and related areas influences thalamic sensory relay activity, providing a potential route for top-down modulation of ascending somatosensory processing ^51^.

The shorter cortical response to tactile stimulation (i.e., N1) observed in Athletes and Artistic swimmers compared with Controls may reflect enhanced functional connectivity among task-related networks in a large set of cortical and subcortical regions involved in sensorimotor processing. Such facilitation has been reported by Giboin et al.^52^ after intensive short slackline training, including the primary and secondary somatosensory cortices, superior parietal cortex, premotor and motor areas, as well as the hippocampus and several cerebellar and brainstem structures. This cortical facilitation likely accounts for the behavioral improvements observed after such a short training period ^53^.

Therefore, the enhanced somatosensory processing observed in dancers and slackliners may result from their intensive training, which repeatedly engages plantar sole tactile afferents to control complex equilibrium states. These findings align with prior work showing that, relative to non-athlete controls, dancers display enhanced N1 SEP amplitude and greater grey matter volume in the S1 representation of the dominant foot, but not the dominant hand^54,55^ (for N1 SEP and S1, respectively). The stronger and shorter tactile-brain coupling observed in Artistic Swimmers points to another explanation. Unlike Dancers and Slackliners, Artistic Swimmers train almost exclusively in water without foot contact with a solid surface. Indeed, their training occurs in an environment where conventional balance equilibrium constraints are mainly absent, and buoyant hydrodynamic forces, rather than contact forces, are responsible for aquatic movements. Therefore, a plausible explanation is that the repeated vestibular stimulation (e.g., head spinning) to which artistic swimmers are exposed, similar to that experienced by dancers and slackliners, may modulate tactile information processing independently of foot use.

Evidence points to a strong functional interaction between vestibular and tactile systems. Ferrè et al.^56^, for example, showed that caloric vestibular stimulation alters tactile detection in the hand: when applied during a skin detection task, vestibular stimulation increases tactile perceptual sensitivity. Because visual sensitivity remained unaltered in the same study, this effect cannot be attributed to a general enhancement of perceptual performance across sensory modalities but instead indicates a selective influence on tactile somatosensory processing. Converging evidence for such sensory interdependence also comes from patients deprived of proprioceptive and tactile inputs, who exhibit enhanced vestibular sensitivity ^57–59^, consistent with compensatory cross-modal reweighting.

Within this framework, the downweighting of the vestibular system, likely resulting from the extensive vestibular stimulation of athletic training ^1,2^, may lead to an upregulation of somatosensory signals to support accurate, tactile-based estimates of body position in space. Support for this interpretation is provided by studies in microgravity, an environment that removes vestibular-induced gravitational inputs, in which plantar tactile stimulation improves perception of body orientation ^10,60^. In this context, foot cutaneous stimulation produces a marked increase in activation of the right posterior parietal cortex ^60^, an important node for vestibular-somatosensory integration and the encoding of whole-body spatial representations ^29^ (for review).

Taken together, these findings suggest that repeated exposure to challenging vestibular stimuli (i.e., fast head motion) can recalibrate multisensory weighting in favour of faster and more effective use of plantar tactile information for balance control. This mechanism, revealed here in athletes, may have direct translational relevance for individuals with partial vestibular loss following concussion, for example, who typically exhibit impaired postural responses and poor recovery following balance perturbations^61^. Systematically exposing these individuals to activities that progressively challenge the vestibular system, such as swings or underwater slackline training, which provides a safer environment, could harness this adaptive cross-modal plasticity to enhance the processing and functional impact of tactile cues. Rather than attempting to restore vestibular function per se, such interventions may promote a compensatory strategy in which enhanced somatosensory-cortical coupling supports rapid and reliable detection of foot-ground interactions and more accurate estimation of body orientation and motion. This perspective opens a rehabilitation avenue focused on driving adaptive sensory reweighting through targeted vestibular stimulation, aiming to improve balance-correcting responses and reduce whole-body instability in patients with vestibular deficits.

## Acknowledgments

We thank Franck Buloup for developing the software Docometre used for data acquisition and Dany Paleressompoulle for developing the set up used for the experiment.

